# Animal model of leprous neuritis using transfer of sensitized lymphocytes into *M. leprae*-inoculated nude mice

**DOI:** 10.64898/2026.07.17.739143

**Authors:** Junichiro En, Masanori Matsuoka, Koichi Suzuki, Masamichi Goto

## Abstract

**Background:** Leprosy is curable with multidrug therapy, but preventing peripheral neuropathy remains challenging. In leprosy, reversal reactions (Type 1 reactions) are characterized by sudden enhanced cell-mediated immunity against *Mycobacterium leprae* (*M. leprae*), leading to acute neuritis that may cause irreversible nerve damage, paralysis, sensory impairment, and muscle atrophy. Armadillo and nude mouse models are used to study *M. leprae* infection, but detailed neuropathological models for immune-mediated reactions in leprosy are lacking. This study aimed to establish a reproducible animal model to analyze leprous neuritis pathogenesis.

**Methodology/Principal Findings:** BALB/c nude mice were inoculated with *M. leprae* and six months later received transfers of naïve CD4^+^ cells, sensitized CD4^+^ cells, or sensitized whole spleen cells from BALB/c mice. Histological and quantitative analyses were performed four weeks post-transfer. Mice that received transfer of sensitized CD4^+^ cells or sensitized spleen cells exhibited footpad swelling up to 88% greater than in animals with no cell transfer. Mice that received the sensitized cells also had massive inflammatory cell infiltration into nerve fascicles. Quantitative imaging confirmed a significant reduction in the percentage of myelinated area in mice that received transfer of sensitized CD4^+^ (p < 0.05) or whole spleen cells (p < 0.01) compared to control mice with no transfer. Furthermore, bacterial fragmentation within the endoneurium coincided with myelinated axon destruction, indicating that the immune response targeted bacilli but simultaneously damaged nerve structures.

**Conclusions/Significance:** This study successfully reproduced leprous peripheral neuritis that resembles the reversal reaction in humans. The findings indicate that CD4^+^ cells are primary drivers of inflammation, whereas interactions between multiple immune cell populations in whole spleen cell transfers further exacerbate disease pathology. This model confirms that host immune responses are crucial for nerve injury progression. This model provides a valuable tool for investigating neuroprotective therapies to prevent permanent disability in patients with leprosy.

**Author Summary:** Although leprosy can be treated as an infectious disease, many patients still suffer from permanent peripheral nerve damage, known as leprous neuritis. A particularly dangerous condition called the "reversal reaction" occurs when the immune system of an infected individual suddenly overreacts, causing severe acute inflammation in the nerves. Without prompt treatment, this inflammatory response can result in irreversible nerve damage, paralysis, and muscle atrophy.

In this study, we developed a new animal model using "nude mice", which lack their own T-cells, to understand how this damage happens. We infected these mice with leprosy bacteria and then transferred specific immune cells, specifically sensitized CD4^+^ T cells, isolated from *M. leprae*-immunized mice. The results showed that this transfer triggered significant footpad swelling and caused inflammatory cells to invade the nerve fascicles. Most importantly, we observed a significant reduction in myelin, the protective coating that promotes nerve signaling.

These findings confirm that nerve damage is largely driven by the host’s own immune response attempting to clear the bacteria. By successfully recreating this reaction, our research provides a vital tool for developing new therapies aimed at preventing permanent disability and protecting the physical functions of patients.

## Introduction

Prevention and treatment of leprous peripheral neuropathy remains a substantial challenge in management of leprosy even after the establishment of multidrug treatments. Several mechanisms are thought to be involved in leprous neuropathy, including inflammation induced by an immune response (leprosy reactions type 1 and 2) [1], damage to Schwann cells via *M. leprae*-specific phenolic glycolipid-1 (PGL-1) without an immune response [2], entrapment neuropathy [3], and disuse atrophy [4]. Among these possible mechanisms, leprosy reaction is the most important in clinical practice. During the course of disease, type 1 reaction (reversal reaction) can occur in which T-cell mediated cellular immunity rapidly increases, causing severe acute inflammation in the skin and peripheral nerves. If left untreated, the reversal reaction can lead to irreversible nerve paralysis and muscle atrophy.

Animal models of leprosy have been established in armadillos and mice. Nine-banded armadillos were experimentally infected with *M. leprae* and their peripheral nerves were examined in a study by Scollard *et al*. [5]. These animals had *M. leprae* within the endothelium of epineurium blood vessels, lymphatics, and intraneural vessels, notably in the absence of accompanying inflammatory lesions.

In mouse models of human lepromatous leprosy, *M. leprae* multiply in the foot pads of athymic nude mice like BALB/c *nu*/*nu* that lack T lymphocytes (T cells) and allow the bacilli to grow systemically [6]. Attempts have been made to induce the reversal reaction using lymphocyte transfer [7], but to date there are no studies on peripheral nerve lesions.

To reproduce the reversal reaction in rodents, Shannon et al. intravenously administered splenic cell suspensions derived from *nu*/+ mice vaccinated with heat-killed *M. leprae* into multibacillary nude mice [8]. Footpad inflammation and swelling, a decrease in bacterial morphological index, and mononuclear cell infiltrations were observed. Notably, however, neuropathological analyses were not performed.

In this study we transferred whole spleen cells or spleen-derived CD4^+^ cells from BALB/c mice sensitized with *M. leprae* into nude mice that were infected with *M. leprae* six months before transfer, and then examined peripheral nerve lesions in the nude mice resulting from a type 1 (reversal) response.

## Materials and Methods

The overall flow of this experiment is shown in Fig 1.

**Fig 1.**
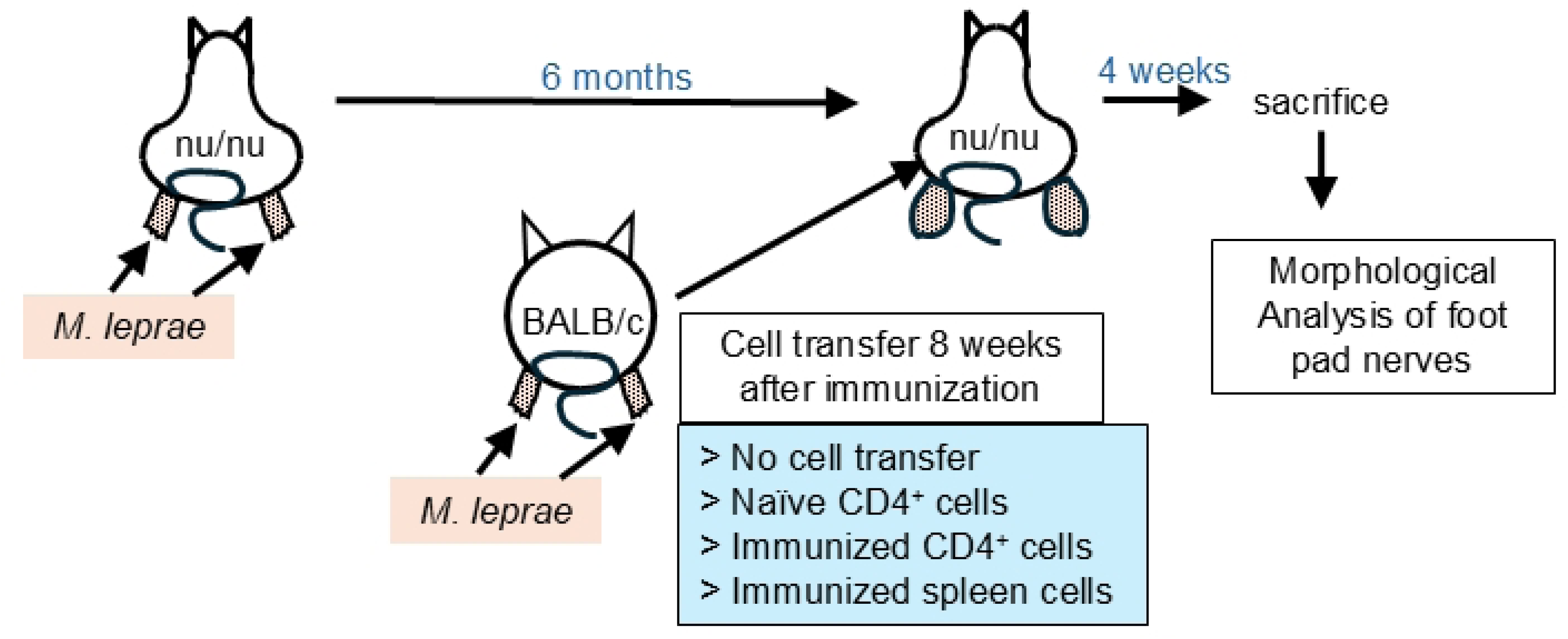
Overall flow of this experiment. BALB/c *nu*/*nu* mice were inoculated with *M. leprae* and the infection was allowed to proceed for six months. Meanwhile, BALB/c mice were immunized by inoculating the sole of the right foot with *M. leprae*. Eight weeks after inoculation, sensitized CD4^+^ cells and spleen cells were isolated. Naïve CD4^+^ cells were isolated from non-immunized BALB/c mice. These different cell types were injected into the peritoneal cavity of BALB/c *nu*/*nu* mice that had been carrying *M. leprae* for six months. There were four experimental groups: no cell transfer, transfer of naïve CD4^+^ cells, transfer of sensitized CD4^+^ cells from immunized mice or transfer of whole spleen cells from immunized mice. Four weeks after the cell transfer, the animals were sacrificed and morphological analyses of foot pad nerves were conducted.

### Inoculation of *M. leprae* into nude mice

A bacillary suspension of *M. leprae* Thai-53 strain, propagated in footpads of nude mice, was prepared by differential centrifugation [9]. A total of 6.0 × 10⁷ bacilli suspended in 50 *μ*L PBS were inoculated into both footpads of BALB/c-*nu*/*nu* mice and allowed to propagate for six months before the transfer experiments.

### Sensitization of normal mice by *M. leprae*

To produce cells for transfer, the bilateral foot pads of BALB/c mice were inoculated with Thai-53 strain *M. leprae* (1.0 x 10^7^). Cells were isolated as described below eight weeks after the inoculation.

### Preparation of lymphocytes and transfer into nude mice

In this study, four experimental groups were used (Fig. 1). **Group A** comprised one *M. leprae*-inoculated mouse that acted as the control and did not receive a cell transfer. **Group B** mice (n=2) received transfer of naïve CD4^+^ cells (n=2). To collect these cells, spleens were first taken from the naïve, non-sensitized BALB/c mice. Lymphocytes were then isolated by Ficoll-Paque (Pharmacia Biotech, U.S.A.) and CD4^+^ cells were collected using magnetic beads (DYNABEADS mouse CD4 (L3T4) and DETACHaBEADS mouse CD4 (Dynal, Norway)) according to the manufacturer’s instructions. Isolated cells (5.0×10^5^) were transferred into the abdominal cavity of *M. leprae*-inoculated nude mice via intraperitoneal injection. **Group C** mice (n=2) received transfer of sensitized CD4^+^ cells. To isolate these cells, spleens were taken from sensitized BALB/c mice and CD4^+^ cells were isolated. The cells (5.0×10^5^) were transferred as described for Group B. **Group D** mice (n=1) received transfer of sensitized spleen cells (5.0×10^5^) isolated from sensitized BALB/c mice as described for Group B.

### Histological examination

Four weeks after cell transfer, all nude mice were subjected to deep anesthesia and sacrificed before fixation with perfusion of 4% paraformaldehyde. At the time this experiment was conducted, an animal experiment committee had not yet been established at the Leprosy Reseach Center’s predecessor, the National Institute for Leprosy Research. Therefore, the animal experiments were carried out with due consideration for the welfare of the laboratory animals. Foot pads were first embedded in paraffin and sectioned into 5 µm-thick sections that were stained using hematoxylin-eosin, Fite’s acid-fast staining. Portions of the foot pads were post-fixed with 1.5% glutaraldehyde and osmium, and embedded in Epon to produce 1 µm semithin sections that were used in electron microscopy examinations.

### Quantitative analysis of light microscopy images of semithin sections

For objective evaluation of peripheral nerve fiber distribution and density, quantitative analyses were performed using light microscopy images of semithin sections. Epon sections (1 µm-thick) were stained with toluidine blue and imaged under strictly standardized optical conditions to ensure comparability across experimental groups. Digital images were acquired using a Nikon FXA light microscope (10× objective, 2.5× internal magnification) equipped with a Fujifilm FinePix S1 Pro camera, capturing at a resolution of 1440 × 960 pixels in JPEG format. Fields for analysis were randomly selected from areas with clearly identifiable nerve bundles to eliminate observer bias.

Image pre-processing was conducted using Photoshop Elements 3.0 (Adobe, U.S.A.). To achieve the highest contrast for peripheral nerve fiber myelin, the red channel was extracted from the RGB palette. The images were converted to grayscale and saved as uncompressed TIFF files to prevent degradation during subsequent analysis.

The processed images were then imported into NIH Image for measurement. The total area of each peripheral nerve bundle was defined by manually tracing the outer boundary using the freehand selection tool. To identify myelinated regions, the "Density Slice" function was employed to apply a standardized threshold corresponding to the intensity levels of the positively stained areas. These thresholds were determined through a preliminary analysis and applied uniformly across all specimens. Multiple fields were measured for each specimen to calculate a representative mean. Finally, the "Positive area ratio (%)" was calculated as: [(Positive area / Total nerve bundle area) × 100] to serve as a standardized index for nerve myelin density.

### Statistical analysis

Quantitative data are presented as the mean ± standard deviation (SD). Comparisons between two groups were conducted using the Mann–Whitney U test. A p value of < 0.05 was considered statistically significant.

## Results

### Foot pad swelling

Initially, the characteristics of the foot pads from mice inoculated with *M. leprae* were examined. First, the area of foot pad cross sections was measured (Table 1). Compared to the control mouse that did not receive a cell transfer, foot pad cross sections from mice that received transfer of naïve CD4^+^ cells were 12% larger, whereas foot pads of mice that received transfer of sensitized CD4^+^ cells and sensitized spleen cells were 56% and 88% larger, respectively.

**Table 1.** Transfer of lymphocytes to *M .leprae*-inoculated nude mice.

| Animal No. | Transferred cells | Cell Type | Foot pad size (mm<sup>2</sup>) | Loss of myelinated fibers from nerves | Cell density in nerves | Bacilli Shape |
| --- | --- | --- | --- | --- | --- | --- |
| NM5-6 | No transfer | na | 12.6 | — | 11.4 | Solid |
| NM5-4 | Naïve | CD4+ | 12.6 | + |  | Solid and granular |
| NM5-5 | Naïve | CD4+ | 15.7 | + | 39.3 | Solid and Granular |
| NM5-1 | Sensitized | CD4+ | 19.6 | ++ | 63.7 | Granular |
| NM5-2 | Sensitized | CD4+ | 19.6 | ++ |  | Granular |
| NM5-3 | Sensitized | Spleen cells | 23.7 | ++ | 64.9 | Granular |

### Foot pad dermal tissue histology

Next, a histological examination of the inoculation site was performed. The control mouse without cell transfer had numerous foamy macrophages that were filled with non-fragmented solid acid-fast bacilli. Focal collection of small numbers of neutrophils was noted. In contrast, mice that received transfer of naïve CD4^+^ cells had decreased numbers of foamy macrophages and a mild increase in the number of small round cells (lymphocytes). A mixture of solid and fragmented acid-fast bacilli was also present. Mice that received transfer of sensitized CD4^+^ cells or sensitized spleen cells showed a remarkable decrease in the number of foamy macrophages and increased density of small round cells. However, no spindle-shaped macrophages or epithelioid cells with a wide cytoplasm were present. Most of the acid-fast bacilli were fragmented or had a granular degenerating pattern.

### Histology of nerves in the foot pad

To determine whether nerve damage was associated with lymphocyte transfer, a careful histological examination of the peripheral nerves was conducted. In control mice that received no cell transfer, macrophages were filled with solid acid-fast bacilli (Fig 2A, left). Meanwhile, in mice that received sensitized CD4^+^ cells, acid-fast bacilli in the endoneurium showed prominent fragmentation or granular degeneration (Fig 2A, right). In all mice examined, about 20% of nerve fascicles contained acid-fast bacilli. In the foot pad from control mice that received no cell transfer, the axons and myelin were well preserved (Fig 2B, left), and a small number of macrophages was observed in the endoneurium. Mice that received transfer of sensitized CD4^+^ cells and those that received sensitized spleen cells both had decreased axonal density and a remarkable increase in the number of small round cells (mostly lymphocytes) (Fig 2B, right). An Epon section (1 µm-thick) from the control mouse had well-preserved nerve fibers (Fig 2C, left), whereas the section from a mouse that received sensitized CD4^+^ cells had marked nerve fiber degeneration (Fig. 2C, right).

**Fig 2.**
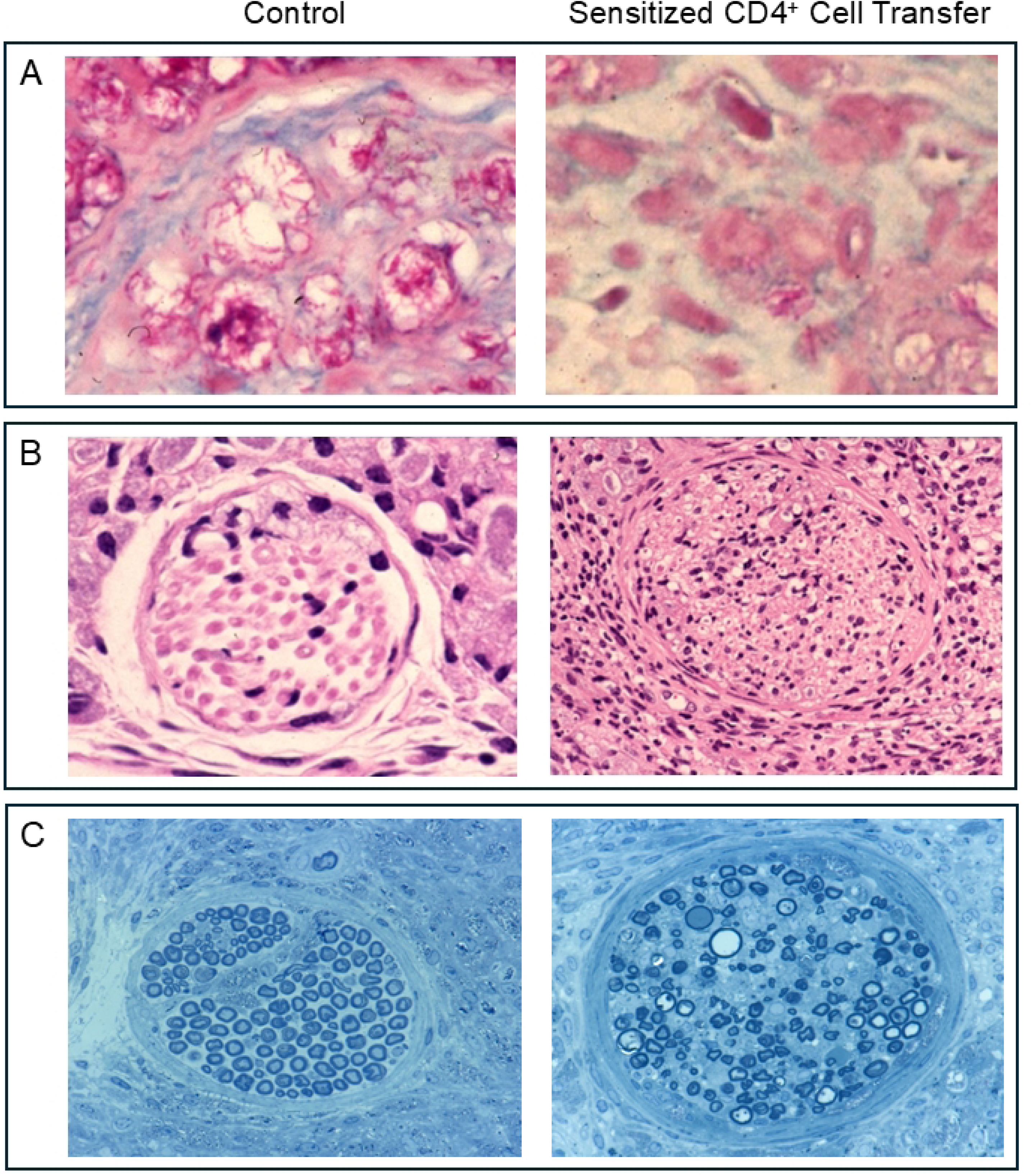
Intradermal nerves in foot pads of *M. leprae*-inoculated nude mice. (A) Acid-fast staining of control mouse (left) shows solid bacilli and mouse with transfer of sensitized CD4^+^ cells (right) shows granularly degenerated bacilli in the nerves. (B) Control mouse (left) shows well-preserved nerve fibers, but a mouse with sensitized CD4^+^ cell transfer (right) shows a decrease in myelinated fibers and inflammatory cell infiltration. (C) Epon sections (1 µm-thick) from the control mouse show well-preserved nerve fibers and a mouse with sensitized CD4^+^ cell transfer (right) showed nerve fiber degeneration.

### Ultrastructure of foot pad nerves

To conduct a more detailed investigation, an electron microscopic analysis was carried out. In the mouse that did not receive cell transfer, the density of myelinated fibers was preserved, and a small number of foamy macrophages filled with solid acid-fast bacilli was present, mainly in the perineurium (Fig 3A). Mice that received transfer of naïve CD4^+^ cells had a mild decrease in the number of myelinated fibers and a small number of macrophages was observed. Mice that received transfer of sensitized CD4^+^ cells (Fig 3B) showed a remarkable decrease in the density of myelinated fibers. Infiltration of lymphocytes, neutrophils and macrophages was also observed in the endoneurium.

**Fig 3.**
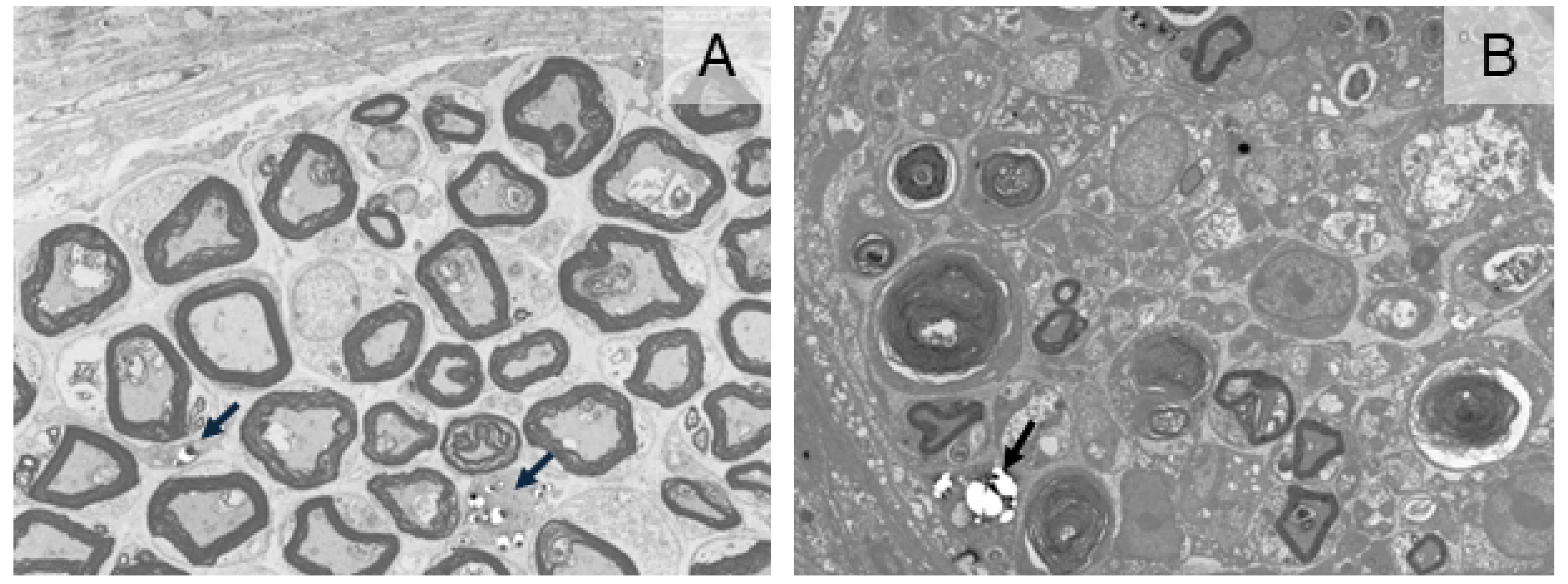
Ultrastructure of nerves in foot pads of *M. leprae*-inoculated nude mice. (A) Control mouse with no cell transfer showed well preserved myelinated nerve fiber density. Foamy macrophages filled with solid acid-fast bacilli were visible (arrows). (B) Ultrastructure analysis of mouse that received transfer of sensitized CD4^+^ cells showed decreased myelinated nerve fiber density as well as reduced lymphocyte infiltration and lower numbers of neutrophils and macrophages.

### Quantitative analysis of light microscopy images

To provide an objective evaluation of nerve fiber density, a quantitative analysis was performed. The positive area (myelin area) within peripheral nerve bundles is summarized in Table 2.

**Table 2.** Mean percentage of positive area in peripheral nerve bundles.

| Experimental group<br>(transferred lymphocytes) | Mean | Standard<br>deviation |
| --- | --- | --- |
| Not done | 18.6% | $\pm 5.6\%$ |
| Naïve CD4<sup>+</sup> cells | 20.6% | $\pm 5.2\%$ |
| Immunized CD4<sup>+</sup> cells | 17.5% | $\pm 4.2\%$ |
| Immunized spleen cells | 10.7% | $\pm 4.2\%$ |

The control group that received no cell transfer showed a mean positive area of 18.6 ± 5.6%. Meanwhile, the group receiving naïve CD4^+^ cells from non-sensitized mice demonstrated an area of 20.6 ± 5.2%, while the group receiving CD4^+^ cells from sensitized mice showed a 17.5 ± 4.2% area, and the group that received transfer of spleen lymphocytes had a mean positive area of 10.7 ± 4.2%.

Figure 4 shows the distribution of the percentage of positive nerve fiber area within peripheral nerve bundles among the different experimental groups. The nerve fiber area did not significantly differ between the group that did not receive a cell transfer and the group that received naïve CD4^+^ cells isolated from non-immunized mice. In contrast, the percentage of positive nerve fiber area was significantly reduced in mice that received sensitized CD4^+^ cells derived from immunized mice (p < 0.05), and an even more pronounced reduction was observed for mice that received transfer of spleen lymphocytes from immunized mice (p < 0.01).

**Fig 4.**
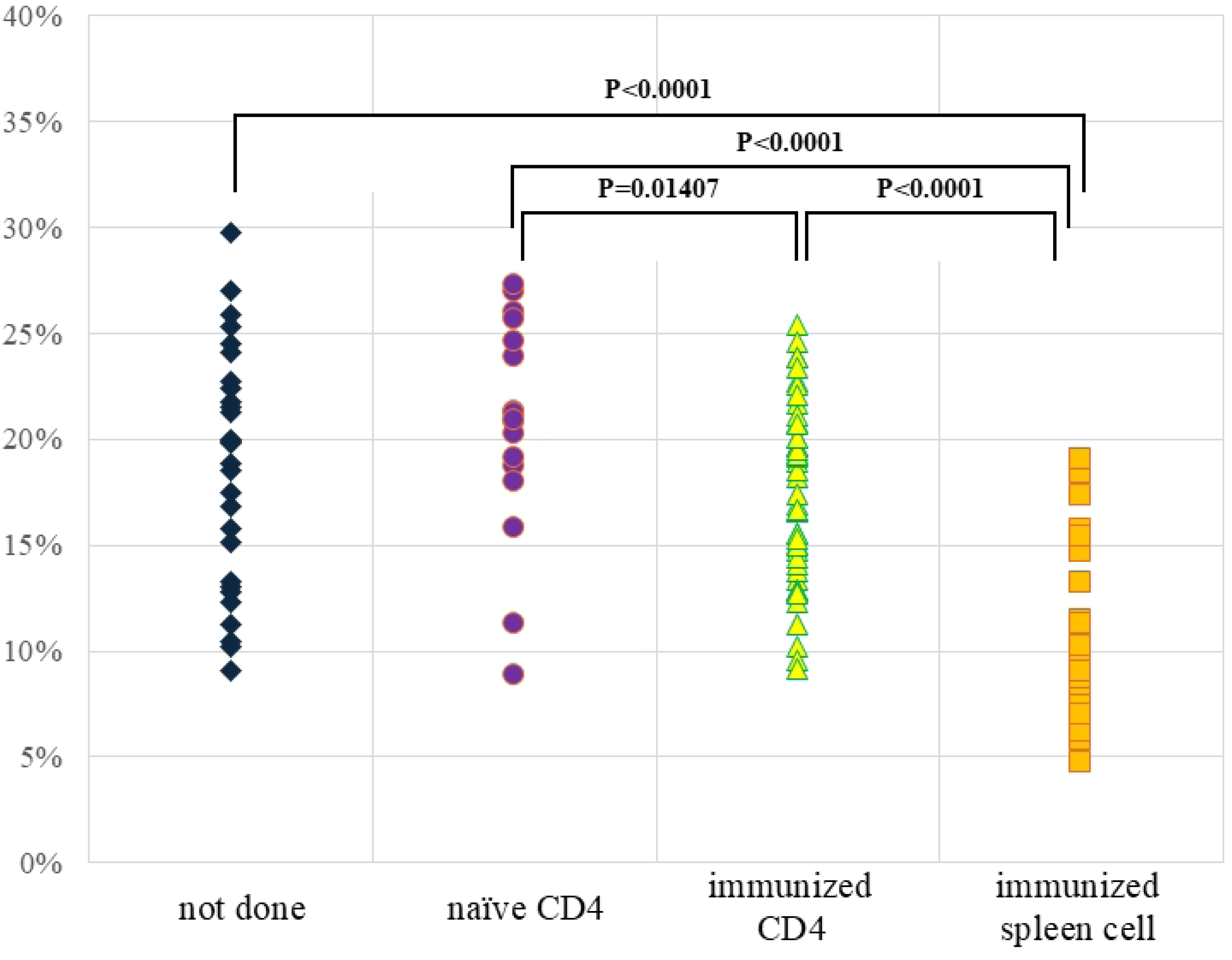
Distribution of the percentage of positive nerve fiber area within peripheral nerve bundles according to the type of lymphocyte transferred. No significant difference in positive nerve fiber area was observed between the control group with no cell transfer and the group that received transfer of naïve CD4^+^ cells derived from non-immunized mice. In contrast, the percentage was significantly reduced in the group that received sensitized CD4^+^ cells derived from immunized mice (p < 0.01), and an even more pronounced reduction was seen for the group that received transfer of spleen lymphocytes isolated from immunized mice (p < 0.0001).

These results indicate that transfer of spleen lymphocytes is associated with a reduction in nerve fibers within peripheral nerve bundles and demonstrates that the severity of peripheral nerve injury varies depending on the type of transferred lymphocyte.

Figure 5 compares the correlation between the area of peripheral nerve bundles and the percentage of positive nerve fiber area among the four experimental groups. In the group with no cell transfer and the groups receiving CD4^+^ cells derived from either non-sensitized or sensitized mice, the percentage of positive nerve fiber area increased as the area of the peripheral nerve bundle increased. Accordingly, the fitted regression curves for these three groups exhibited an upward slope, indicating that nerve fibers were relatively preserved in larger peripheral nerve bundles.

**Fig 5.**
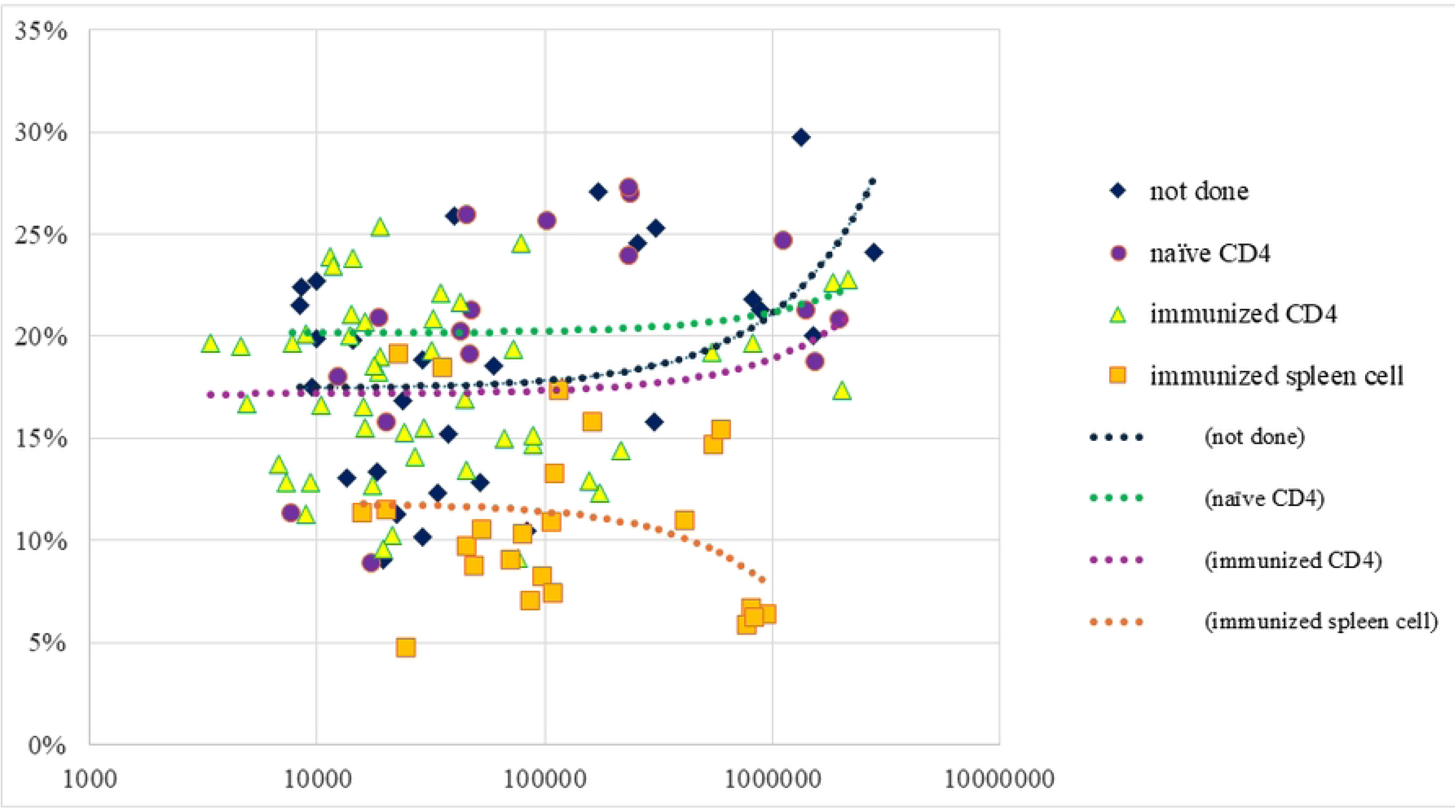
Correlation between the area of peripheral nerve bundles and the percentage of positive nerve fiber area among the four experimental groups. For the control group that received no cell transfer and the groups receiving CD4^+^ cells derived from either non-immunized (naïve) or immunized (sensitized) mice, the percentage of positive nerve fiber area increased as the area of the peripheral nerve bundle increased. These data indicate that nerve fibers were relatively preserved in larger peripheral nerve bundles. Meanwhile, mice that received transfer of spleen lymphocytes isolated from immunized mice had a reduced percentage of positive nerve fiber area even in peripheral nerve bundles that had larger areas, indicating that nerve fiber damage also occurred in larger nerve bundles in this group.

In contrast, the group that received transfer of spleen lymphocytes had a reduced percentage of positive nerve fiber area even in peripheral nerve bundles that had larger areas. As a result, the fitted regression curve for this group showed a slightly downward slope, that differed from the trends seen for the other three groups. This finding indicates that nerve fiber damage occurred not only in smaller peripheral nerve bundles but also in larger ones in the group that received transfer of spleen lymphocytes.

Collectively, these results demonstrate that the relationship between peripheral nerve bundle size and nerve fiber preservation varies according to the lymphocyte type transferred, indicating that the different transfer groups had distinct patterns of peripheral nerve injury.

## Discussion

Animal models of the reversal reaction in leprosy involving thymus transplantation [7] or transfer of spleen cells [8] into *M. leprae*-inoculated nude mice have been reported, but to our knowledge there are not yet detailed studies of nerve lesions in these reversal reaction models.

In this study, we investigated whether leprous peripheral neuritis could be induced in *M. leprae*-infected nude mice. To explore this possibility, we conducted experiments using four groups of BALB/c nude mice in which *M. leprae* was allowed to proceed for six months. These groups had no lymphocyte transfer (control), transfer of CD4^+^ cells isolated from non-immunized (naïve CD4^+^ cells) or immunized (sensitized CD4^+^ cells) BALB/c mice, or transfer of spleen lymphocytes isolated from immunized BALB/c mice. The mice that received CD4^+^ cells or spleen cells from immunized mice exhibited increased footpad swelling as well as inflammatory cell infiltration into nerve fascicles, reduced density of myelinated nerve fibers, and fragmentation of acid-fast bacilli. These findings are considered to reflect immune-mediated neuropathy associated with the type 1 reaction, also termed reversal reaction, in leprosy.

Peripheral neuropathy in leprosy is thought to result not only from the affinity of *M. leprae* for peripheral nerves, particularly Schwann cells, but also from host immune responses that exacerbate neural damage. Spierings et al. [10] reported that, during reversal reactions, *M. leprae*-infected Schwann cells can act as antigen presenting cells. Interactions with inflammatory T cells may subsequently induce Schwann cell injury, leading to peripheral nerve damage. In particular, cell-mediated immune responses centering on CD4^+^ cells are considered to play a critical role in the pathogenesis of leprous neuropathy.

In the present study, the BALB/c nude mouse that received no lymphocyte transfer exhibited foamy macrophages containing acid-fast bacilli within nerves, but the axonal and myelin structures were largely preserved. These findings suggest that the mere presence of *M. leprae* is not sufficient to induce severe inflammatory neuropathy, but rather that host immune responses are essential for nerve injury progression. Moreover, BALB/c nude mice that received transfer of CD4^+^ cells isolated from non-immunized mice had mild lymphocyte infiltration and minimal neural degeneration. In contrast, the BALB/c nude mice that received transfer of CD4^+^ cells isolated from immunized mice showed marked lymphocytic infiltration into nerves, decreased axonal density, and myelin degeneration. These findings suggest that antigen-specific sensitized CD4^+^ cells reacted against *M. leprae*-infected neural tissues and induced inflammatory neuropathy. Similar neural lesions were also observed in mice that received transfer of spleen lymphocytes isolated from immunized mice and this group had the most pronounced reduction in the ratio of myelinated fiber-positive area. Splenic lymphocytes contain not only CD4^+^ cells but also various immune cell populations, including CD8^+^ cells and B cells. Therefore, mice with transfer of spleen lymphocytes exhibited not only direct effects of CD4^+^ cells, but interactions among multiple immune cell populations may have amplified inflammatory responses, resulting in more extensive nerve damage. Indeed, neural injury in leprosy is suggested to involve interactions among T cells, macrophages, Schwann cells, and *M. leprae* [10], and the present findings support these previous observations.

Morphological changes in acid-fast bacilli were also an important finding. In the mouse that received no lymphocyte transfer, solid bacilli predominated, but in mice that received sensitized CD4^+^ cells or spleen lymphocytes transferred from immunized mice, fragmented bacilli and granular degeneration were frequently observed. These findings suggest that the transferred immune cells activated local macrophages and promoted bactericidal activity against *M. leprae*. This immune response contributed to bacterial clearance, but also simultaneously induced damage to peripheral nerve tissues. Thus, this experimental model appears to reproduce the pathological condition observed in clinical reversal reactions, in which “immune responses against bacilli” and “neural tissue injury” progress simultaneously.

Neural injury caused by *M. leprae* is thought to involve not only immune-mediated mechanisms but also direct pathogenic effects that are independent of immune responses. PGL-1, a cell wall component specific to *M. leprae*, is reportedly involved in Schwann cell binding and neurotropism. Ng et al. demonstrated that PGL-1 contributes to *M. leprae* binding to peripheral nerves [11]. More recent studies also suggested that PGL-1 may induce metabolic alterations and neurotoxic phenotypes in Schwann cells [12]. Therefore, leprous neuropathy is considered to develop along with Schwann cell injury associated with the *M. leprae* neurotropism with CD4^+^ cell-mediated immune responses further exacerbating inflammatory neural damage.

One notable finding of the present study was that mice that received sensitized CD4^+^ cells or spleen lymphocytes isolated from immunized mice had reduced density of myelinated fibers, but the nerve fascicles were not completely destroyed, and some neural fiber structures were preserved. Quantitative analyses also demonstrated reduced, but not completely absent, myelinated fiber-positive area suggesting that residual nerve fibers persisted even after inflammatory neural injury.

In conclusion, the present study demonstrated that transfer of sensitized CD4^+^ cells or whole spleen cells from immunized mice into *M. leprae*-infected nude mice successfully reproduced leprous peripheral neuritis that resembles the reversal reaction. CD4^+^ cells appear to play a central role in the formation of neural injury, whereas multiple immune cell populations contained within whole spleen cells may further exacerbate reversal reaction pathology.

### Study limitations

After initial inoculation of nude mice, leprosy bacilli require 6 months to multiply locally. Therefore, in this study, the main experiment was performed only once after investigating the optimal conditions (data not shown). This small study sample raises concerns about reproducibility, although the fact that similar results were obtained in multiple mice suggests that the experimental system is indeed valid.

## Data availability

Raw data were generated at the Leprosy Research Center, National Institute of Infectious Diseases and National Leprosarium Hoshizuka-Keiaien. Derived data supporting the findings of this study are available from the corresponding author [JE] on request.

## Acknowledgments

The authors thank Ms. Taki Hatanaka (Research Laboratory of National Leprosarium Hoshizuka-Keiaien, Kagoshima, Japan) for her excellent technical assistance.

